# A How-To guide to: Quantitative high-speed video profiling to discriminate between variants of primary ciliary dyskinesia

**DOI:** 10.1101/614966

**Authors:** M. Chioccioli, L. Feriani, Q. Nguyen, J. Kotar, S. D. Dell, V. Mennella, I. Amirav, P. Cicuta

**Affiliations:** Biological and Soft Systems Sector, Cavendish Laboratory, University of Cambridge, Cambridge, UK; Pulmonary, Critical Care and Sleep Medicine Section, Dep. Internal Medicine, Yale University School of Medicine, New Haven, CT, USA; Institute of Clinical Sciences, Imperial College London, London, UK; Department of Biochemistry, University of Toronto, Toronto, Canada; Department of Pediatrics, SickKids Hospital, University of Toronto, Toronto, Canada; Cell Biology Program, SickKids Hospital, Toronto, Canada; Department of Pediatrics Respirology, University of Alberta, Edmonton, Canada

## Abstract

This document presents detailed steps to analysing the waveforms of beating cilia, measured in high speed video microscopy. We show that in the case of PCD caused by mutations in *DNAH11* and *HYDIN* the classification by a handful of parameters describing ciliary dynamics allows to distinguish the genotype, as well as (much more easily) distinguishing healthy from PCD samples. This document is intended to complement the brief highlight (1) and presents the details of the datasets used in that letter.

## 1. Materials and methods

### 1.1 Cell isolation

In the study (1), human primary nasal airway cell samples from healthy subjects and patients with PCD caused by mutations in *DNAH11* and *HYDIN* (see Table 1 for details), were collected from University of Alberta Department of Pediatrics, SickKids Hospital, Toronto Canada and NJH Denver CO, USA.

**Table 1.** Subjects with primary ciliary dyskinesia used in this study (*siblings). SD1b, SD2b and SD1, SD2 were from the same donors but cells grown at ALI and videos recorded at different times.

| Subject | Gene | Mutations | Sites | N cilia |
| --- | --- | --- | --- | --- |
| A1 | HYDIN | c.2616_2617insTGGCACTGAC; p.Leu873TrpfsTer3 | Univ. of Alberta | 4 |
| A2 | HYDIN | c.2616_2617insTGGCACTGAC; p.Leu873TrpfsTer3 + c.14428-1_14431delGGTGA | Univ. of Alberta | 3 |
| A3* | DNAH11 | c.9239T>C, p.Leu3080Pro | Univ. of Alberta | 2 |
| A4* | DNAH11 | c.9239T>C, p.Leu3080Pro | Univ. of Alberta | 2 |
| A5 | DNAH11 | c.6041-1G>T + c.8870C>G, p.Ala2957Gly. | Univ. of Alberta | 6 |
| A6 | DNAH11 | c.13516_13522dupGTTCTGG, p.(Ala4508Glyfs*14); c.4942C>T, p.(Gln1648*) | Univ. of Alberta | 7 |
| SD1 | DNAH11 | Cys4296Stop; Ile4122Ser | SickKids | 10 |
| SD1b | DNAH11 | Cys4296Stop; Ile4122Ser | SickKids | 12 |
| SD2 | HYDIN | c.10368-2A>G listed as a canonical splice site; (c.8674C>G;8675delA) (p.Gln2892Glyfs*3) frameshift deletion with adjacent substitution | SickKids | 14 |
| SD2b | HYDIN | c.10368-2A>G listed as a canonical splice site; (c.8674C>G;8675delA) (p.Gln2892Glyfs*3) frameshift deletion with adjacent substitution | SickKids | 13 |
| SD3 | Healthy |  | SickKids | 6 |
| MS2 | Healthy |  | NJH | 2 |
| MS3 | Healthy |  | NJH | 4 |

University of Alberta: human cells were collected with a cytology brush (Celletta™ brush cell collector with protective tip product number 9100060; Engelbrecht Medizin-und Labortechnik GmbH, Tiefenbachweg 13; 34295 Edermünde, Germany), as part of routine clinical investigation. To minimize pain, the brush was rinsed with an isotonic saline solution before being inserted in the inferior nasal meatus, using rotatory and linear movements. The brush was immediately removed and transferred into a tube containing 2–3 mL of cell culture medium (RPMI Medium 1640, Endotoxin tested, Cell culture tested, without L-Glutamine, without Sodium Bicarbonate. Biological Industries LTD., Beit Haemek, Israel). The tube was then shaken vigorously so that cells became detached from the brush.

SickKids, University of Toronto: Cells from patients at the SickKids Hospital in Toronto were collected with a cytology brush as described above and plated on collagen-coated (PurCol, Bovine Collagen Type I, Advanced BioMatrix) T-25 flasks (Corning) with PneumaCult-Ex Medium (Stem Cell Technologies) supplemented with ROCK Inhibitor (5 μM; Y-27632; Enzo Life Sciences) and hydrocortisol solution (96 ng/ml; Stem Cell Technologies). Under these conditions, basal (stem) cells were then expanded, seeded on transwells (Corning HTS Transwell-96 and -24 permeable support; 0.4 μm pore size), and differentiated following Stem Cell Technologies protocols using PneumaCult-Ex and PneumaCult-ALI media at 37°C with 5% CO_2_. This differentiation process spanned over a minimum 35-days period and was repeated at two different period of time: SD1, SD2 cells were grown at ALI and HSVM performed in 2017; for SD1b, while SD2b cells were grown at ALI and HSVM performed in 2018. PneumaCult-Ex and -ALI media were supplemented with the following antibiotics: Vancomycin (100 μg/ml; Bioshop Canada Inc), Tobramycin (80 μg/ml; Bioshop Canada Inc), Gentamicin (50 μg/ml Thermo Fisher Scientific) and Antibiotic-Antimycotic (1 X; Thermo Fisher Scientific).

University of Cambridge, Prof Pietro Cicuta’s lab: Human primary nasal airway cell samples from two healthy donors were kindly provided by Prof. Max Seibold (National Jewish Hospital, Denver, CO, USA). The healthy cells were collected, expanded and maintained as previously described (2). Briefly, Cells from patients at NJH were collected with a cytology brush as described above and plated on Culture vessels coated with Rat Tail Collagen, Type I, 30µg/ml in PBS without Ca^++^/Mg^++^ (Cat#354236, BD Biosciences, San Jose, CA). At P0 nasal cells were cultured in BEGM plus singlequots (Lonza, Basel, Switzerland). Once cells reached P1 all BEGM media components including antibiotics and antifungals were added as directed. Cells were cultured until ∼80% confluency. Passage 4 cultured cells were differentiated using the previously described method (2). Briefly, cells were plated at a density of 90,909 cell/cm^2^ (12mm semipermeable inserts) in ALI expansion medium containing. At confluence, the expansion medium was switched to ALI differentiation media. ALI culture was maintained for 28 days.

### 1.2 Recording HSVM

Samples in the study (1) were observed via high speed video microscopy using the following set-ups:

University of Alberta: approximately 50 microliters of the cell suspension was used to perform HSVM: cells were imaged under an inverted microscope, Eclipse TS-100 inverted phase contrast microscope (Nikon Corp. Tokyo, Japan), with a 40X objective lens. Videos were recorded at 120 frames per seconds (fps) for 3 seconds, using a scA640-120 fm digital video camera (Basler AG, Ahrensburg, Germany) (3).

SickKids University of Toronto: cilia from each sample type were imaged by placing the filters cell-side down on a 1.5cm round coverslip mounted on a holder. HSVM (500fps; 8s videos) was performed using spinning disc confocal microscopy. The protocol was approved by Sickkids Ethics Committee.

University of Cambridge: cells at ALI were shipped to the Cicuta laboratory (University of Cambridge, United Kingdom). To perform HSVM on the cilia, we first excised the bottom membrane – supporting the epithelium - from the plastic ALI insert. The membrane was then folded in half and placed between two coverslips, together with a few μl of medium. The profile videos were then obtained by imaging the edge of the folded membrane. High speed videos were acquired at room temperature on an Eclipse Ti-E inverted microscope (Nikon, Japan) using a 60X water immersion objective (NA = 1.2, Nikon, Japan) and an additional 1.5X magnification in the optical path, and a CMOS camera (GS3-U3-23S6M-C, FLIR Integrated Imaging Solutions Inc., Canada). The image acquisition was run by a custom software running on a Linux machine. About 10 videos were acquired per insert, in different locations along the membrane’s edge. Videos were at least 10s long, recorded with a framerate in excess of 200fps (1px = 0.065 μm).

### 1.3 Quantitative measurements on the ciliary beating pattern

The HSVM videos have to be analysed to recover cilia positions in successive frames. This can be done manually using an image editor and storing coordinates of pixels on one cilium. In the study (1) this process was carried out in Prof Cicuta’s lab in Cambridge using a new user interface developed in Matlab (The MathWorks). The interface was designed to help the user to manually tracking the cilium position over time. Once the cilium positions on one frame are stored, this is repeated for each frame. This allows to extract the time-evolution of the evolving shape, which can be smoothed using an appropriate mathematical function.

Based on these shapes, we determined the <u>amplitude and velocity</u> of the ciliary beat at various distances (2, 3, 4 and 5 μm) from the cell surface (Figure 1-A). The <u>curvature</u> at every point along the cilium could then be determined, and the <u>average curvature</u> across different tracked cilia was expressed as a function of the relative position along the cilium length (Figure 1-B), Knowing the displacements of the cilium across subsequent frames, t<u>he force exerted by the different parts of the cilium onto the surrounding fluid</u> was calculated using Resistive Force Theory, following the method described in (4) (Figure 1-C).

**Figure 1.**
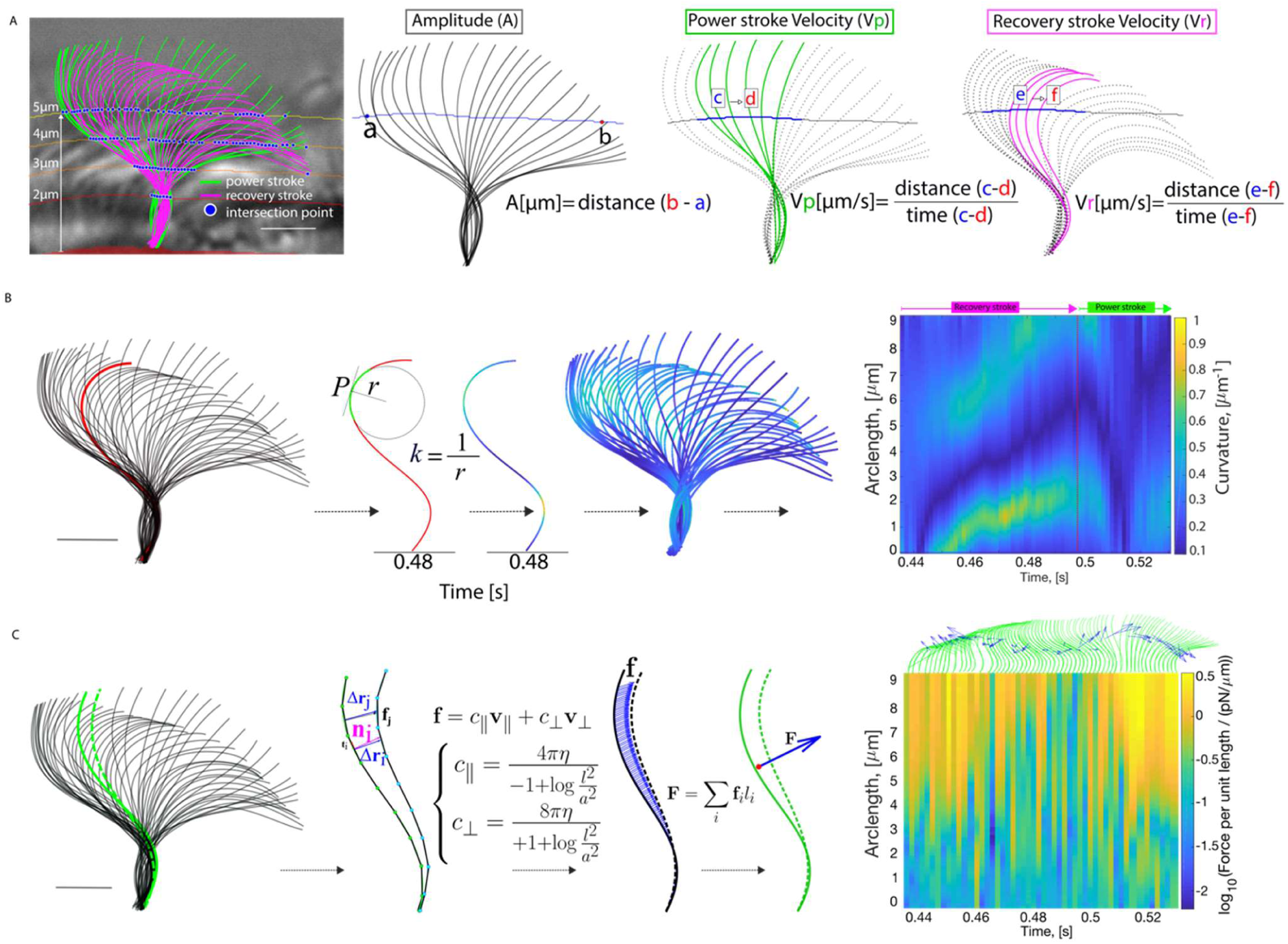
(A) Obtaining the amplitude and velocity of the ciliary beat. Cilium beating amplitude and velocity during the power and recovery stroke can be measured at any given distance from the cell surface by drawing a line that intersects all individual tracks at a given distance from the cell surface. The amplitude is measured as the distance between the intersection points of the cilium with each measuring line that are further apart from each other. The velocity is measured by dividing the distance between subsequent intersection points by the time interval. (B) Measuring the curvature of the cilium over time. The waveform extracted from the tracking algorithm allows curvature (κ) to be measured at every point along the cilium (see supplementary methods for equations). The cilium shape is then coloured according to its local curvature. Local values for curvature (showed in colours) are used to compose a kymograph, where the evolution of the curvature can be tracked. Note the pattern of high-low-high curvature (green - dark blue - green) that propagates from the base to the extremity of the cilium during the recovery stroke. The power stroke shows instead a lower curvature overall, as the cilium moves in an extended configuration. The waveform shown in this figure is reconstructed from a healthy control. Scale bar is 2µm. (C) Calculating the force exerted by the cilium on the fluid during the ciliary beat cycle. Resistive force theory is applied to the data extracted from tracking individual cilia (in this case healthy control) to calculate the force per unit length exerted by the cilium on the surrounding fluid during the evolution of the beat cycle (4); (see methods for further details and equations). Local values for the force density can then be built into a kymograph that shows the magnitude of the force per unit length (colour) as a function of the position along the cilium (y axis) and of time (x axis).

The details of the tracking and interpolating of the cilium shape over time are given in the following sections.

#### 1.3.1 Tracking and reconstruction of the ciliary beating pattern

In each pre-recorded frame, the user clicks P points along the cilium shape (with, on average, P = 7), obtaining P couples of coordinates (X_i_, Y_i_), with i being an index that grows from 1 to P from the base to the tip of the cilium. These points need not be exactly evenly spaced. A smooth representation of the cilium is then reconstructed by fitting the coordinates of the clicked points with two 4^th^ degree polynomials. First, we measure the Euclidean distance between subsequent couples of points: 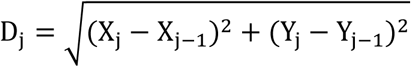. We then define the parameter S_i_, corresponding to the coordinates (X_i_, Y_i_), as the sum of all the Euclidean distances between the base and the i-th point: 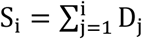, with S_1_: = 0. The discrete series of data points [(S_1_, X_1_), … (S_P_, X_P_)] and [(S_1_, Y_1_), … (S_P_, Y_P_)] are finally fitted with two 4th degrees polynomials A and B, so that 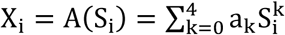 (and likewise Y_i_). A much smoother representation of the cilium can then be obtained by evaluating the polynomials on s, a finely spaced variable ranging from 0 (at the base of the cilium) to S_P_ (at the tip of the cilium), resulting for each frame in a parametric representation of the cilium as (x(s), y(s)), where 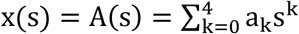 and 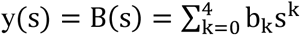.

#### 1.3.2 Finding the measuring lines at a constant distance from the cell surface

Still in the custom user interface the user creates a binary mask that closely follows the cell surface. Example of such masks can be seen in Figure 1-A and 2 The measuring line, for example at a distance *h* from the cell surface, is found by performing a dilation of the binary mask using a disk of radius *h* and taking the edge of the dilated mask.

#### 1.3.3 Measuring the amplitude and the cilium velocity during power and recovery stroke

The amplitude of the cilium beat at distance h from the cell surface is measured as the distance between the two furthermost points where the cilium would intercept the measuring line at height h (“intersection points”) (Figure 1-A). By taking the intersection points of the cilium in two subsequent frames and measuring the distance between them, and dividing it by the time interval, we measure the instantaneous values of velocity (Figure 1-A). We can then average all velocities measured on displacements towards the power stroke direction (“power stroke velocity”) and, separately, towards the recovery stroke direction (“recovery stroke velocity”).

#### 1.3.4 Measuring the curvature

In each frame, the unsigned curvature of the cilium described by (x(s), y(s)) is measured as 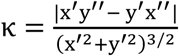, where the derivatives are in the parameters (Figures 1-B, 3). To average the curvature across different tracked cilia we express the curvature as a function of the relative length of the cilium, a variable that ranges from 0 (base of the cilium) to 1 (Figure 1-B). As the cilium beats its tip moves in and out the focal plane, and is therefore hard to track consistently. To normalise the cilia length, we discard very short regions of some tips, assigning the value “1” to the length of the reconstructed cilium that was the shortest during the beating cycle. Cilia shape could not be determined in some few frames, but we were able to retain analysis of the video as a whole by reconstructing the cilium shape in the missing frame by interpolating the coefficients of the two 4^th^ degrees polynomials (x = A(s), y = B(s)) between the previous and subsequent frames, i.e. assuming that the shape of the cilium changes smoothly over time (Figure 1-B). To address the possibility that using 4^th^ degrees polynomials to reconstruct a smooth mathematical model for the cilium shape might lead to overestimating the curvature at the proximal and distal end of the cilium, we tracked and reconstructed nine immotile cilia, and measured their curvature along the length of the cilium. We then subtracted the curve obtained averaging across the immotile cilia from each of the active cilia’s curvature curves.

#### 1.3.5 Measuring the force

Resistive Force Theory (RFT) can be used to measure the force exerted by the cilium on the fluid based on the method described in Brumley et al., 2014 (4). To measure the force exerted in the time between two frames, first we approximated the reconstructed snapshot in each of the two frames with a series of cylinders about the direction parallel and perpendicular to the cylinder itself. We can then define the velocity the cylinder moved at as the ratio between its displacement and the time interval between the two frames (Figure 1-C). If we define the velocity of the i-the cylinder as v_i_, and its two components as (v_i⊥_, v_i‖_), then thanks to RFT we can find the force a cylinder exerts on the fluid as f_i_ = (c_⊥_v_i⊥_, c_‖_v_i‖_)l_i_. l_i_ is the length of the i-the cylinder, and 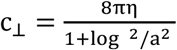 and 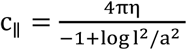 have the units of viscosities and are the drag coefficients of a slender cylinder of length l and radius a. The total force is then found by integrating the force contributions of all the cylinders, considering overlapping cylinders. The position of the centre of drag in a frame is finally obtained by performing an average over the position of all the cylinders’ midpoints, weighted with their contribution to the total force (Figure 1-C).

### 1.4 Statistical analyses

Results in (1) are presented in the figures as mean ± SD unless otherwise specified in the caption. First, we calculate the average across tracked cilia from the same patient, then mean and SD are calculated across all patients within the same group (healthy, *HYDIN*, or *DNAH11*). The SD is indicated in the figures either with error bars (Figure 4-B) or as shaded regions (Fig. 4S-A, C-E, G, 5S). To average several curves [y_1_ = f_1_ (x), y_2_ = f_2_(x), …, y_N_ = f_N_(x)] measured each on a different tracked cilium, for each value x* of the independent variable we averaged the values of the dependent variable 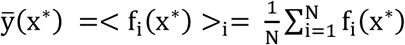. At the same x*, the standard deviation is measured as 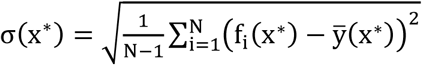. Statistical analysis was performed using unpaired Student’s t-test (with the option for unequal variance) using MATLAB. The 95% confidence level was considered significant.

### 1.5 Principal Component Analysis (PCA)

We ran Principal Component Analysis (PCA), the results of which are in Figure 2 of ref. (1), on a subset of N = 85 well-characterised cilia. The N cilia are the subset of all analysed cilia in which all of the measurements detailed in this work were undertaken. The subset was composed of cilia from 4 *HYDIN* patients, 5 *DNAH11* patients, and 3 healthy patients. The five specific parameters to build the “barcode” for our samples were, in order: i) the amplitude of the beat at 5μm from the cell surface, ii) the time averaged curvature measured at mid-length, iii) the standard deviation of the curvature over time measured at mid-length, iv) the autocorrelation of the curvature, measured at Δs=0.15, and v) the mean force exerted on the fluid, measured at nine tenths of the cilium length. The first three Principal Components (PCs) calculated on the N=85 by d=5 dataset account for 88% of the total variance of the sample. In the plane defined by the first two PCs the data points representing the average of the cilia from the same patient form two non-intersecting clusters, with data from the healthy samples very well separated from the PCD samples. An unsupervised clustering algorithm (k-means by MATLAB, 15 repetitions) separated data from healthy and PCD samples in 2 different clusters, only based on the projection of the data along the first 2 PCs. The data points of the two PCD variants then form two separated, non-intersecting clusters in the plane defined by the second and third principal component, according to their genotype.

**Figure 2.**
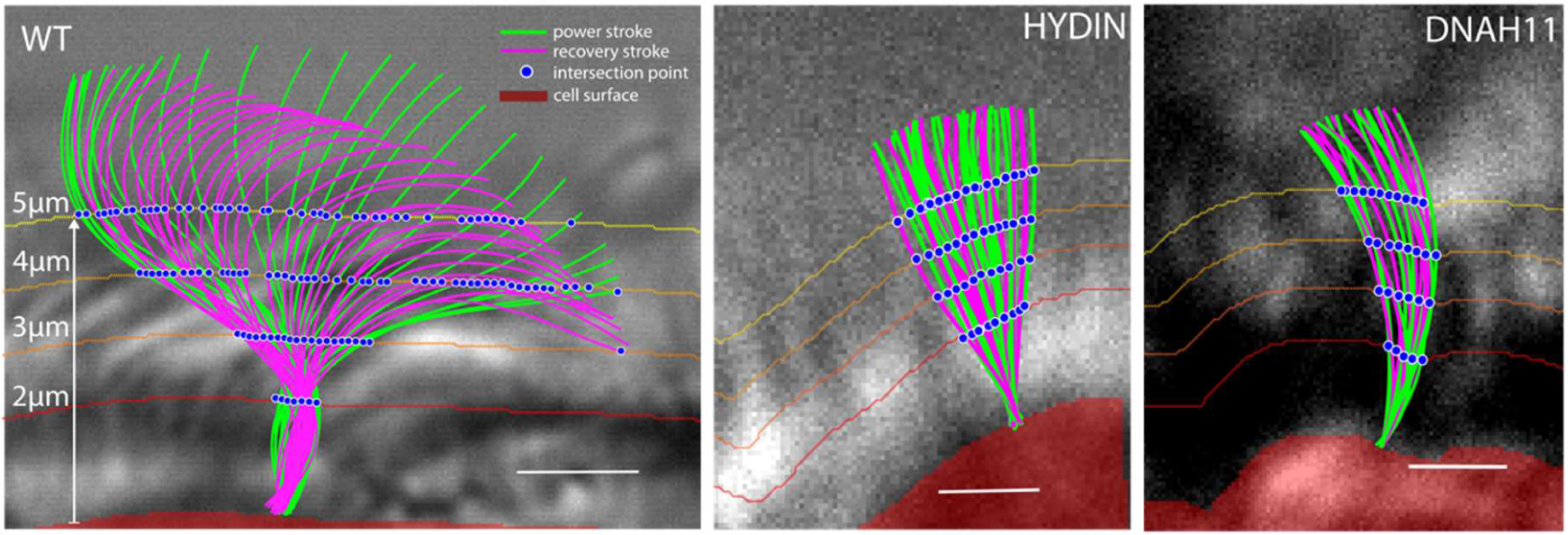
The amplitude is measured as the distance between the intersection points of the cilium with each measuring line that are further apart from each other. The velocity is measured by dividing the distance between subsequent intersection points by the time interval.

## 2. Results and observations

### 2.1 Dynamics and waveforms of ciliary beating from the HSVM analysis pipeline

Once the HSVM data are analysed to extract the “barcode” of amplitude, velocity, force, and curvature of the cilium, it is apparent that there are strikingly differences between the three conditions. Some of these differences would be qualitatively clear by inspection of the waveforms, the most obvious being the severely restricted beating amplitude and curvature of the cilium in cells with mutations in either *DNAH11* or *HYDIN*, compared to healthy cells (Figure 1 of ref. (1), Figure 2). This is consistent with qualitative data previously reported for these mutations (5–7). However, the reconstructed waveforms for the *HYDIN* and *DNAH11* mutant HAECs are also more subtly distinct from each other: qualitatively one could see that the beating amplitude of the *DNAH11* mutant cilia is slightly reduced compared to that of *HYDIN*, and the curvature appears to be more pronounced. Our pipeline allows the waveform dynamics to be interpreted more quantitatively, which we expect to be key for extracting biophysical parameters that drive mucociliary clearance (8), and to gain understanding of how these parameters are affected in PCD. In order to obtain these insights, further data processing is required.

### 2.2 Principal component analysis discriminates between DNAH11 and HYDIN PCD variants

Our HSVM analysis allows us to quantify several properties of the cilium shape. In particular we have focussed on measuring the beat amplitude, the average velocity of the cilium during its power and recovery strokes, the curvature of the cilium throughout the beat along its length, and the amount of force the cilium exerts on the surrounding fluid in a beat cycle. While the following section of this work will describe these quantities in detail, here we show how even a subset of these is sufficient to build a quantitative “barcode” unique to each patient mutation, and that this barcode can discriminate between different PCD variants. We applied Principal Component Analysis (PCA) to the set of all the analysed cilia in which all of the measurements mentioned above were undertaken, choosing just five specific parameters to build “barcodes” for our samples: i) the amplitude of the beat, ii) the curvature of the cilium, measured at mid-length and averaged for the time cycle, iii) the standard deviation of the curvature over time measured at mid-length, iv) the autocorrelation of the curvature and v) the mean force exerted on the fluid.

Briefly, PCA is a standard technique to find new, uncorrelated observable quantities to describe the dataset, starting from a set of possibly-correlated measured quantities. The new observables are called Principal Components, and are linear combinations of the original measured quantities. PCA is often used to reduce the number of variables needed to describe the dataset and can identify the few most uncorrelated variables.

In Figure 2 of ref. (1) (left), we show how the data of the first two Principal Components (P.C.) cluster, so that the healthy and PCD data points are perfectly separated. This separation in the space of the first two P.C., between the data from healthy and PCD patients, was confirmed by an unsupervised k-means clustering algorithm, which successfully separated data from healthy and PCD samples based exclusively on the quantitative barcode of the phenotype, and no previous knowledge of the genotype. While the data points of the two PCD variants do not form separated, non-intersecting clusters in the plane of the first two P.C., they do so in the space of the first, second, and third principal components (Figure 2 of ref. (1)). This shows how PCA on a comprehensive set of measured quantities derived from HSVM (the “barcode”) can be used to discriminate between cells from healthy subjects and those with PCD caused by mutations in *DNAH11* and *HYDIN*.

### 2.3 Amplitude and velocity of the cilium during the beat cycle

The amplitude of the beating cilia at 5 μm from the apical surface was 1.3 ± 0.6 μm and 2.2 ± 1.1 μm for the *DNAH11* and *HYDIN* mutant forms respectively, compared to 8.7 ± 1.6 μm in the healthy control (Figure 4-A). The beat amplitude of the cilia increased approximately linearly with the distance from the cell surface, although the gradient was markedly reduced in the PCD samples as compared to healthy control. These quantitative data are consistent with descriptions of reduced ciliary beating amplitude previously reported for both the *DNAH11*- and *HYDIN*-mutant forms of PCD (6, 7).

**Figure 3.**
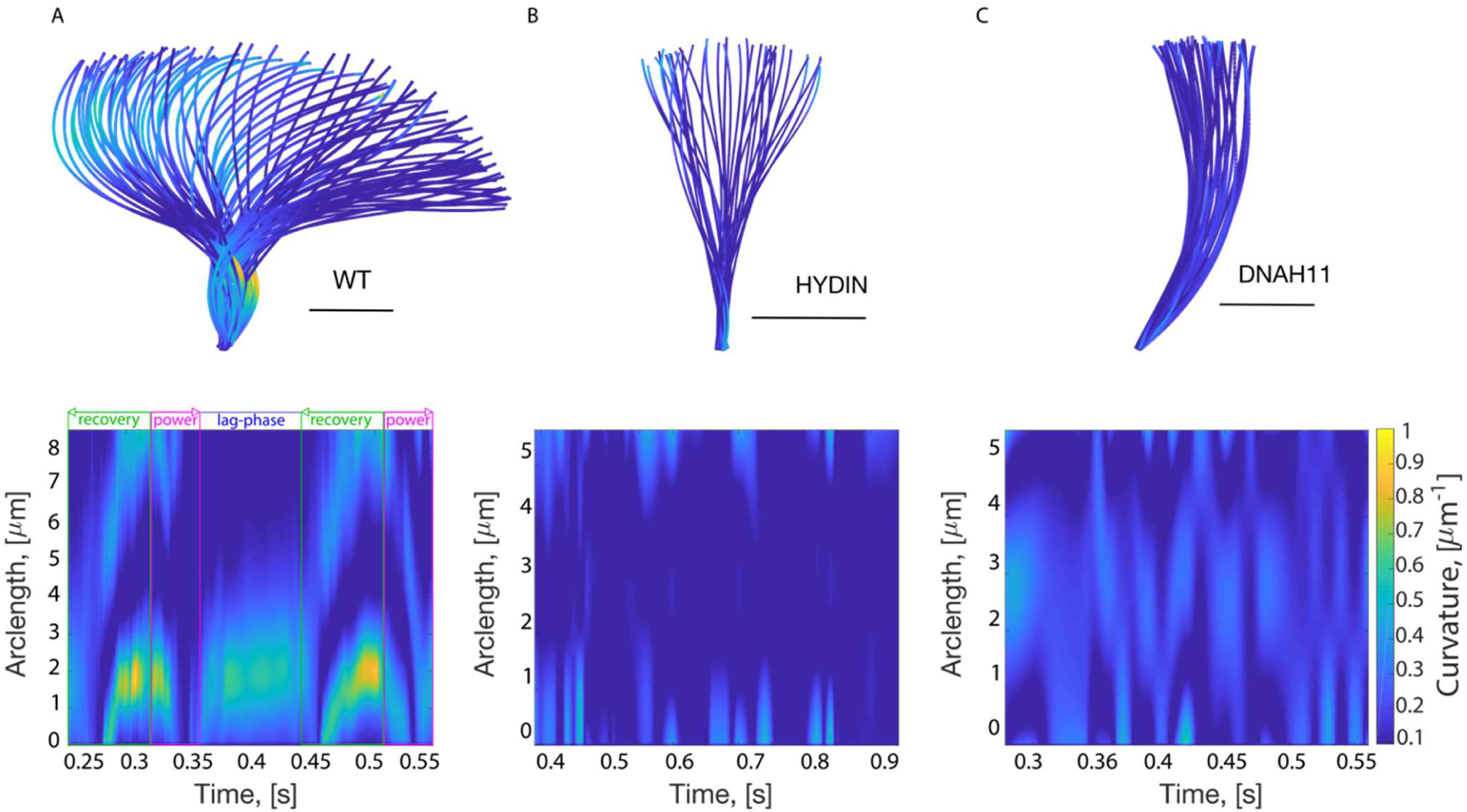
Representative curvature kymographs of single cilia in healthy control and PCD samples highlight very different dynamics of curvature along the cilia. Reconstructed cilium waveforms (top) and heat-maps (bottom) show the time evolution of the curvature of a single representative cilium in healthy (A), HYDIN (B) and DNAH11 (C) samples. Strongly marked and persistent diagonal signals are only present in the heathy cilia. Scale bar is 2μm.

**Figure 4.**
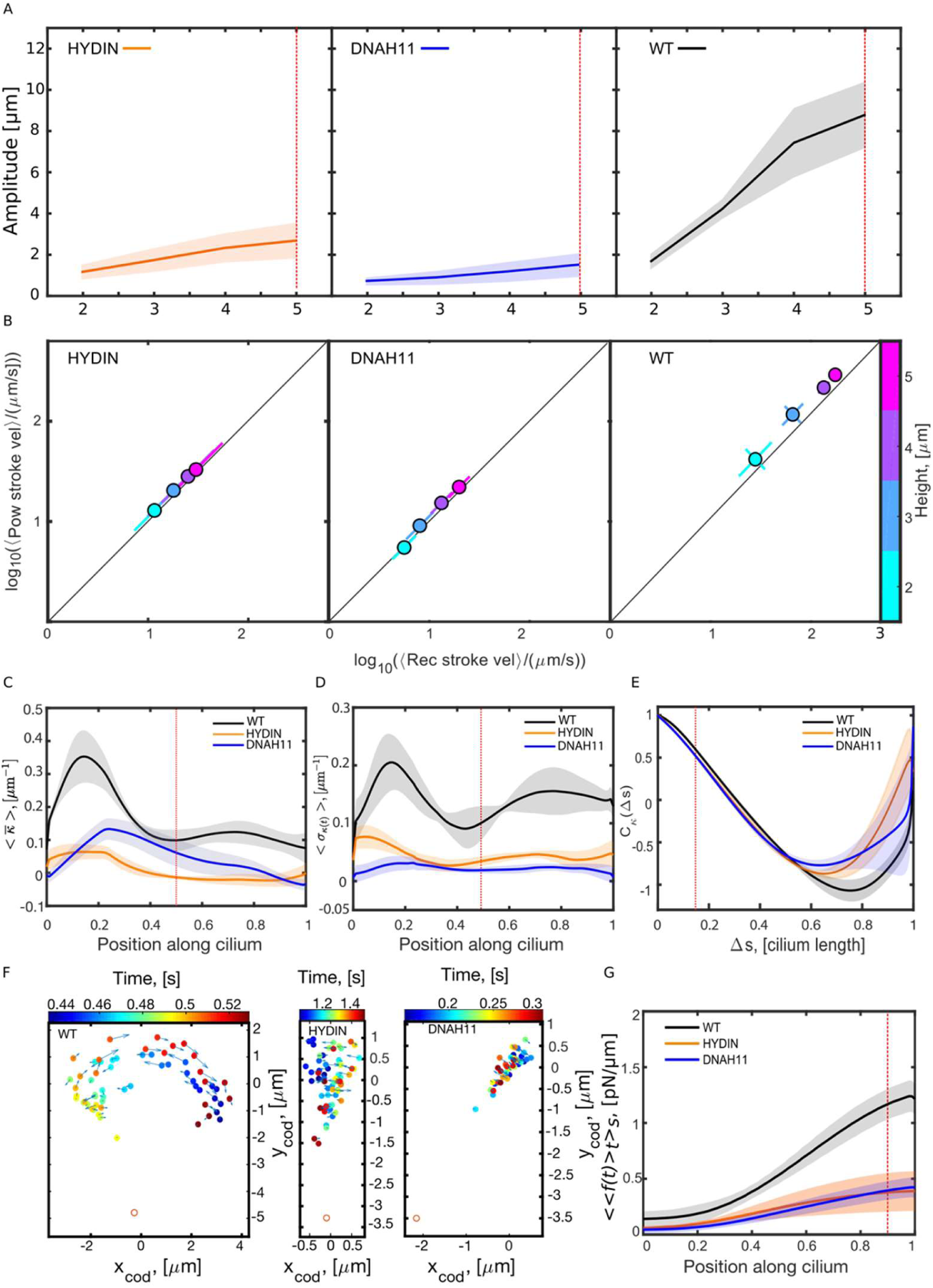
Parameters of the cilium beating pattern. (A) The average amplitude of beating cilia, measured as in Figure 1S and 2S, increases at increasing distances for healthy control, HYDIN and DNAH11 variant PCD forms, although at different rates. (B) Average power stroke velocity vs average recovery stroke velocity at increasing distance for healthy control, HYDIN and DNAH11 variant PCD forms. Colour codes the height from the cell surface. By plotting the velocity of the power stroke on the y-axis, and of the recovery stroke on the x-axis, it becomes easy to see both a difference between the two velocities (points depart from the y=x line, on the side of the higher velocity) and a general increase in the total velocity of the cilium (lower velocities will be shown as data points closer to the origin of the axes, higher velocities will be shown further from the origin). Here we plot the population average taken across data points at the same height from the cell surface (markers). The error bars show the standard deviation in the direction parallel and perpendicular to the y=x line (drawn in black). This was done because the points averaged formed an elongated cluster oriented along the y=x line (Fig. 7S). (C) Time-averaged curvature relative to cilia length, plotted as population average (solid lines) and SD (shaded region). These curves are significantly different: P values calculated with an unpaired two-tailed Student’s t-test at mid-cilium are < 0. 05 between each pair of curves. (D) Standard deviation of curvature over time relative to cilia length, plotted as population average (solid lines) and SD (shaded region). The Healthy curve is significantly different from the two PCD curves: P values calculated with an unpaired two-tailed Student’s t-test at mid-cilium are < 0. 05 between Healthy and HYDIN, and < 0. 01 between Healthy and DNAH11. (E) Time-averaged autocorrelation of the cilium’s curvature, plotted as population average (solid lines) and SD (shaded region). The Healthy curve is significantly different from the two PCD curves (P value calculated with an unpaired Student’s t-test at Lis = 0. 15 was < 0. 05 between the Healthy and each of the two PCD curves). (F) The cilium can be coarse-grained as a point moving along the trajectory defined by the cilium’s centre of drag over time. (G) Time-averaged magnitude of the force density along the cilium in healthy control and DNAH11 and HYDIN PCD variant forms, plotted as population average (solid lines) and SD (shaded regions). The red dotted lines throughout the figure highlight the subset of parameters used to build the “barcode” for Principal Component Analysis.

The velocity of the beating cilium increased with the distance from the cell surface, and was lower in PCD cells compared to healthy cells (Figure 4-B). We next considered whether there was any difference in velocity between the power and recovery strokes and, if so, whether this was constant along the cilium. Whereas healthy beating cilia exhibited a higher average velocity of the power stroke relative to the recovery stroke, this was not the case in the PCD samples. In the *DNAH11* mutant multiciliated cells, the average velocity of the power stroke was higher only at measurements most distal from the cell surface (4 μm and 5 μm), whereas at the proximal end there was no measurable difference (Figure 4-B). In the *HYDIN* mutant cells, there was no detectable difference between the average velocity of the power and recovery stroke at any distance along the cilia. Thus, while there is a distinct difference in the velocity of the power and recovery strokes in healthy beating cells, this difference is lost in cilia from PCD patients with mutations in either *DNAH11* or *HYDIN*.

### 2.4 Curvature of the cilium evolves along the arc length and during the beat cycle

A kymograph of curvature from the base of the cilium to its tip and throughout the beat cycle shows very obvious distinct spatio-temporal patterns among the different samples, indicating distinct travelling waves of curvature along the cilium (Figure 1-B). We generated kymographs for the curvature of the beating cilium over one or more beat cycles in different cilia of HAECs obtained from healthy subjects and subjects with mutations in *DNAH11* and *HYDIN* (Figure 3). In contrast to healthy beating cells, there was no distinct pattern of dynamic curvature in the beating cilia from cells with mutations in either *DNAH11* or *HYDIN* (Figure 3-B-C). Representative kymographs from individual cilia show that the overall curvature in both cases was low, reaching a maximum of 0.4 μm^-1^ and 0.7 μm^-1^ for *DNAH11* and *HYDIN* respectively. In the case of *HYDIN*, curvature was extremely low in the central region of the cilium (2 μm to 4 μm distal to the base, Figure 3-B. This was not observed in the cells obtained from subjects with mutations in *DNAH11*; in this case, all regions of the cilium display a consistently low degree of curvature along the entire cilium at various time points during the beat cycle (Figure 3-C). Finally, no pausing between the end of the power stroke and the beginning of the recovery stroke was detected based on the analysis of the curvature of beating cilia from cells obtained from subjects with mutations in *DNAH11* or *HYDIN*, unlike in healthy samples (Figure 3-A). The kymographs displayed in Figure 3 each represent the curvature of a single beating cilium from each genetic background (healthy, *DNAH11* and *HYDIN*). In order to strengthen our analysis, we applied the analysis pipeline on different cells imaged in multiple different locations to obtain a measure of the average curvature, the standard deviation of the curvature, and the autocorrelation of the cilia curvature over time across the entire sample population (Figure 4, C – E). In the samples from PCD patients, the average curvature was much lower than the one from healthy cells, and appeared lowest overall in the *HYDIN* mutant form (Figure 4-C). Interestingly, in cells from PCD caused by mutations in *DNAH11*, average curvature was much higher in the central region of the cilia than at the proximal and distal ends. The degree to which the curvature changes at any given position along the cilium throughout the beating cycle can be measured by taking its standard deviation. The population-averaged standard deviation of the curvature for PCD samples caused by mutations in both *DNAH11* and *HYDIN* were both lower than the one observed in healthy cells, and did not vary as markedly along the arc length of the cilium. This was consistent with the reconstructed waveforms of individually tracked cilia obtained from each genetic background (Figure 3). The autocorrelation of the curvature along the cilium arclength measures how rapidly changes in curvature happen along the cilium (Figure 4-E). Here, the curve obtained from the healthy samples shows a slightly slower decay than both the *DNAH11* and *HYDIN* variant PCD samples.

### 2.5 Force exerted by the cilium during the beat cycle

We calculated the force per unit length exerted by a cilium on the surrounding fluid throughout a beat cycle using Resistive Force Theory, as described in section 1.3.5. Integrating the force density along the cilium we can measure the direction and magnitude of the total force during the beating cycle (Figure 1-C).

Examining the kymograph of the force density throughout the beat cycle in a representative healthy sample (Figure 1-C) we easily observe the power and recovery strokes, which correspond to the two areas of high force represented in the heat-map, and the transition between them (the area with low force density as shown by the kymograph, and where the total force changes direction). In contrast to the behaviour of healthy beating cilia, the force produced by cilia of cells with mutations in either *DNAH11* or *HYDIN* is minimal, and it not obvious to identify the power and recovery stroke solely based on the force density kymographs (Figure 6). We next mapped the position of the “centre of drag” of the cilium over time, by calculating the average of the cylinders’ positions weighted with the modulus of the force they exert on the fluid (Figure 1-C, 4-F). In healthy beating cilia, the centre of drag shows a wide orbital motion throughout the beat cycle, whereas in both the PCD variants the centre of drag remains clustered in a small region, consistent with the reduced amplitude of the beat. The motion of the centre of drag has been shown (8, 9) to be important in synchronising cilia dynamics to enable mucociliary clearance.

**Figure 5.**
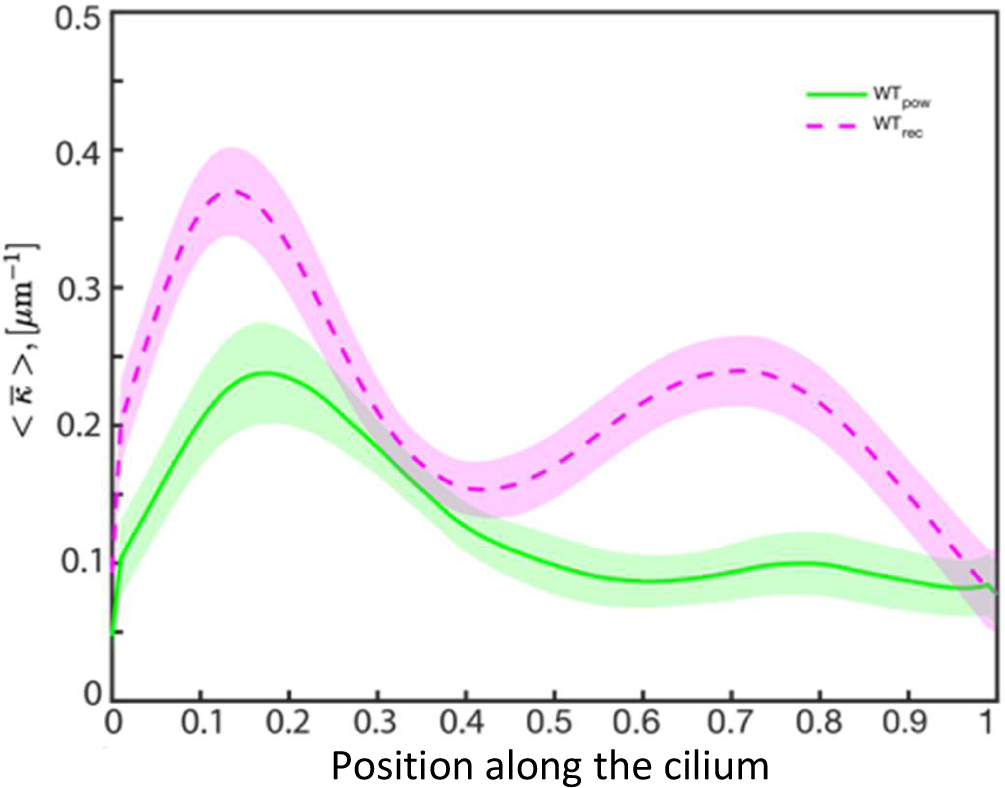
Average curvature of healthy cilia during power and recovery stroke.

**Figure 6.**
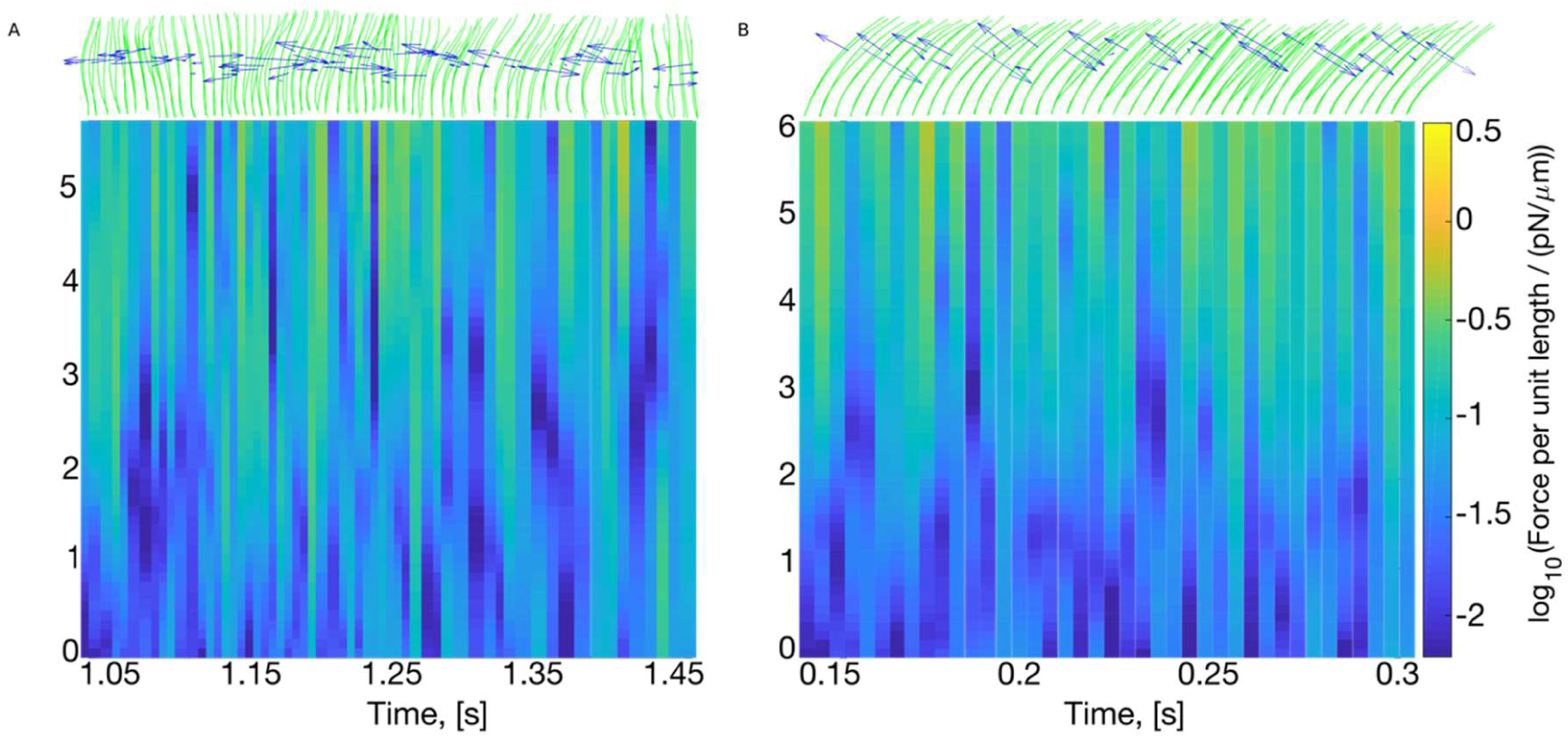
Representative kymographs show the magnitude of the force per unit length (colour) as a function of the position along the cilium (y axis) and of time (x axis) in the HYDIN and DNAH11 PCD variant forms.

**Figure 7.**
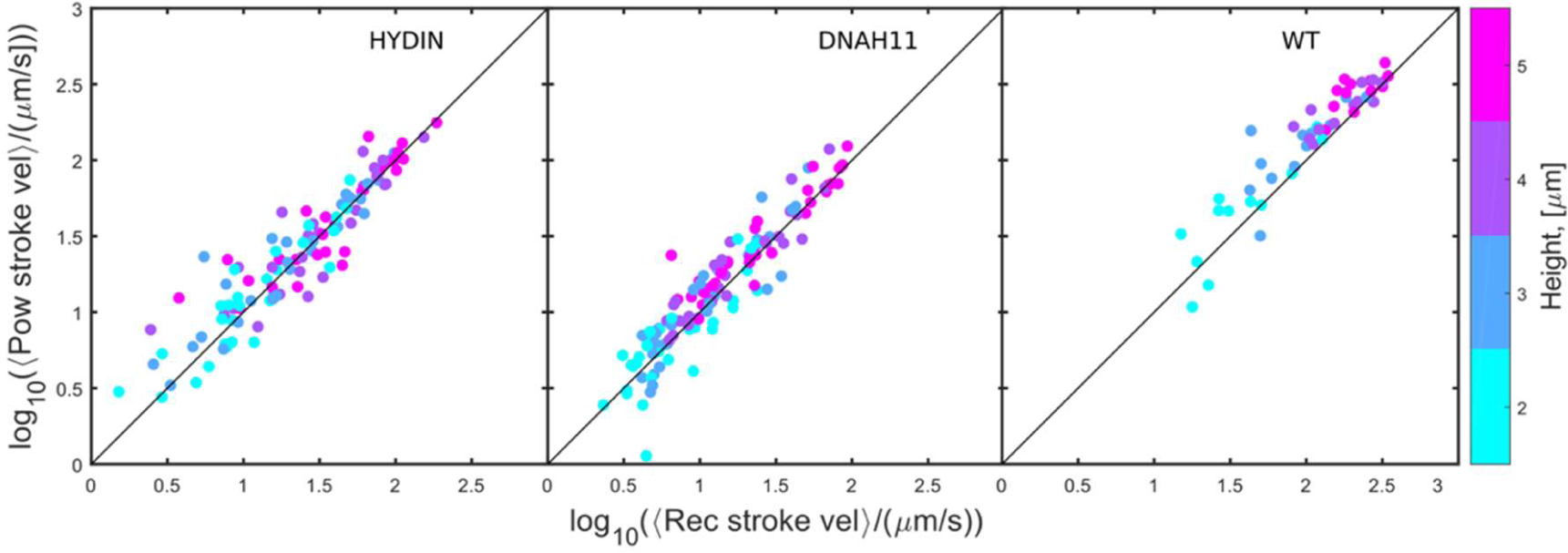
Scatterplots of the average power stroke velocity vs average recovery stroke velocity at increasing distance for healthy control, HYDIN and DNAH11 variant PCD forms.

In all samples, the force per unit length of the cilium increases from base to tip (Figure 4-G), consistent with previous reports (10). This is expected: as the distance from the base of the cilium increases the corresponding point on the cilium must cover increasingly larger amplitudes in the same amount of time. In the population of healthy beating cilia, the force exerted at the distal end is approximately three as large as the force exerted by cilia from cells with mutations in *DNAH11* and in *HYDIN*. Consistent with the data from single cilia (Figure 6), the distribution of force increases gradually for cells with mutations in *HYDIN*, while for *DNAH11* mutant cells the force increases gradually, but only starting from approximately the most distal two-thirds of the cilium arc length: the most proximal third remains constant.

## 3. Considerations on the data and method

We have presented here, and highlighted in (1), an analysis strategy that enables an in-depth characterisation of ciliary beating defects observed in PCD, which in turn allows us to assign unique quantitative profiles to specific PCD variants with mutations in either *DNAH11* or *HYDIN* and to distinguish between them. Our analysis is based only on standard HSVM data as input, and does not require the use of specialised microscopy or experienced clinicians. We anticipate that a quantitative approach to characterising ciliary beating defects will advance both the diagnostic and mechanistic understanding of PCD, particularly in cases where no ultrastructural defect or causal mutation is detected.

Recent progress in the physics of cilia synchronisation has highlighted the importance of both the amplitude of a single beating cilium as well as its velocity over the cycle in collective dynamics and hence mucociliary clearance (11). In current practice, however, these two characteristics are not typically extracted from the HSVM data. Instead, the ciliary beat frequency (CBF), which is the number of full ciliary beating cycles per second, is evaluated for diagnostic and mechanistic studies (15). However, the CBF is not the key parameter behind the physics of ciliary waves (11), and neither is it an informative metric for the diagnosis and analysis of variant forms of PCD, or even PCD versus non-PCD (12–15). For example in patients with the *HYDIN* variant form of PCD, the CBF is reportedly normal (5, 16) and thus indistinguishable from healthy cells. In our new analysis, in contrast, the *HYDIN* variant PCD cilia were clearly distinguishable from healthy control on the basis of velocity and amplitude. Previous attempts to quantify velocity and amplitude of ciliary beating have been reported (9, 16) but these studies analysed only the movement of the tip of the cilia relative to the base, and thus did not consider the dynamic nature of each parameter along the entire arc length of the cilium nor variations across the beat cycle. In our study, we show that these data are absolutely required to build an accurate reconstruction of the ciliary waveform and to quantify important differences between PCD *DNAH11* and *HYDIN* variants.

Many cases of PCD are characterised by defects in ciliary bending (5, 16): both the *HYDIN*- and *DNAH11*-causing PCD variants are known to show an impaired beat pattern, and have respectively been described as “bending capacity reduced” (6) and “reduced bending/increased stiffness” (5). Clearly, however, this does little to distinguish between the two forms. Quantitative analysis of the dynamic curvature of *DNAH11* and *HYDIN* variant PCD cilia revealed a unique profile for each variant form and localised differences in curvature according to the relative proximal/distal position along the cilium. Our principal component analysis shows how, taken together, these unique profiles constitute a quantitative barcode that can be used to discriminate between different variant forms of PCD, and between PCD and healthy samples. Our analysis of individual parameters allows some interesting observations, for example we observe how for the *DNAH11* genotype, both the force and the average velocity of the power stroke relative to the recovery stroke are perturbed only at measurements most distal from the cell surface, and not at the proximal end. This is intriguing, as the proximal region of the axoneme is where the *DNAH11*-driven motors are located, and yet this is not where the effect is observed. This suggests that the downstream effect of *DNAH11* protein activity is more distal than its actual localization. Further work should therefore focus not only on the proteins that physically interact with *DNAH11*, but also those that may act more distally.

We envisage that the quantitative analysis of cilia beat phenotypes presented in this study could be applied to analyse all variant forms of PCD. These data would be used to establish a reference database of ciliary beat “barcodes”: a quantitative dataset for the dynamics of motile cilia, containing the ciliary beat amplitude, velocity, curvature and force, unique to each PCD variant form. Diagnostic implementation of this proof-of-principle study would depend on the collection of a much larger dataset spanning the entire range of PCD-causing mutations, which could then be used as a training set for a machine learning algorithm. Many laboratories already use HSVM as a primary method to diagnose PCD, however the evaluation criteria vary between groups and the data are largely qualitative. Furthermore, current HSVM methods give little insight into the nature of the beating defect itself, where it manifests on the cilium, when it occurs during the beat cycle. A quantitative database of ciliary beat barcodes unique to each PCD variant would be useful in aiding diagnosis, particularly in cases where no ultrastructural defect or known causal mutation is detected, and would further aid in unravelling the genetic basis of PCD.

## Acknowledgments

### Support for the study

This work was supported by EU ERC grant CoG HydroSync (M.C., L.F., J.K., P.C.), V.M. received funding from the Canadian Lung Association.

### Author contributions

M.C. designed the overall study. M.C. and L.F. performed experiments, interpreted data, generated figures and drafted the text. L.F. developed the analysis software. J.K. designed and optimized the HSVM apparatus and acquisition software. P.C. obtained funding, interpreted data and provided overall steering and support. I. A. and Q.N. performed cell culture and collected HSVM movies. M.C, L.F., I. A., P.C., V.M. and S.D. edited the manuscript. We thank C. Hendry for rigorous and thorough manuscript edits, J. Avolio for help with nasal cell scrapings, A. Fitzpatrick for help with HSVM acquisition, and M.A. Seibold for healthy control samples. Competing interests: The authors declare that they have no competing interests. Data and materials availability: software and digital videos data used in this study are available upon request. We thank C.E.Hendry for rigorous and thorough manuscript edits, J.Avolio for help with nasal cell scraping. A.Fitzpatrick for help with HSVM movies acquisition, and M.A.Seibold (NJH, Denver, CO, USA) for kindly providing healthy control samples.

